# Liftoff: an accurate gene annotation mapping tool

**DOI:** 10.1101/2020.06.24.169680

**Authors:** Alaina Shumate, Steven L. Salzberg

## Abstract

Improvements in DNA sequencing technology and computational methods have led to a substantial increase in the creation of high-quality genome assemblies of many species. To understand the biology of these genomes, annotation of gene features and other functional elements is essential; however for most species, only the reference genome is well-annotated. One strategy to annotate new or improved genome assemblies is to map or ‘lift over’ the genes from a previously-annotated reference genome. Here we describe Liftoff, a new genome annotation lift-over tool capable of mapping genes between two assemblies of the same or closely-related species. Liftoff aligns genes from a reference genome to a target genome and finds the mapping that maximizes sequence identity while preserving the structure of each exon, transcript, and gene. We show that Liftoff can accurately map 99.9% of genes between two versions of the human reference genome with an average sequence identity >99.9%. We also show that Liftoff can map genes across species by successfully lifting over 98.4% of human protein-coding genes to a chimpanzee genome assembly with 98.7% sequence identity.

**Availability:** The source code for Liftoff is available at https://github.com/agshumate/Liftoff

## Introduction

Recent developments in DNA sequencing technology have greatly reduced the time and money needed to sequence and assemble new genomes. Currently there are 13,420 eukaryotic genome assemblies in GenBank, of which ~10,000 have been added in the last 5 years alone. The addition of new and improved genome assemblies is a starting point for genetic studies of many species; however, to be maximally useful, the genes and other functional elements need to be annotated. Unfortunately, the annotation of new genomes has not kept pace with sequencing and assembly. This is evident in GenBank, where only 3,540 of the 13,420 eukaryotic genomes have any annotation at all. Eukaryotic genome annotation is a challenging, imperfect process that requires a combination of computational predictions, experimental validation, and manual curation. Rather than repeating this costly process for each new genome that is assembled, a more scalable approach is to take the annotation from a previously-annotated member of the same or closely-related species, and then map or ‘lift over’ gene models from the annotated genome onto the new assembly. In addition, for well-studied organisms, multiple assemblies may be produced over time, and there is an ongoing need to lift the annotation onto these newer, more contiguous assemblies. The most well-known example of this is the human genome, but other model organisms such as mouse, zebrafish (Church *et al.*, 2011), rhesus macaque (He *et al.*, 2019), maize (Jiao *et al.*, 2017), and many others had a series of gradually improved assemblies.

Current strategies for this task use tools such as UCSC liftOver (Kuhn *et al.*, 2013) or CrossMap (Zhao *et al.*, 2014) to convert the coordinates of genomic features between assemblies; however, these tools only work with a limited number of species and they rely only on sequence homology to find a one-to-one mapping between genomic coordinates in the reference and coordinates in the target. This strategy is often inadequate when converting genomic intervals, like a gene feature, rather than a single coordinate. If the interval is no longer continuous in the target genome, current strategies will either split the interval and map it to different locations, or map the spanned interval to the target genome (Gao *et al.*, 2018). In many cases, this disrupts the biological integrity of the genomic feature; for example, if the interval is split and mapped to different chromosomes or strands, or spans a large genomic distance, it may not be possible for it to represent a single gene feature. Furthermore, prior tools convert each feature independently, so while every exon from one transcript may be lifted over to a continuous interval, the combination of exons in the target genome may not necessarily form a biologically meaningful transcript. Mapping each feature independently also often results in multiple paralogous genes incorrectly mapping to a single locus.

Here we introduce Liftoff, an accurate tool that maps annotations described in General Feature Format (GFF) or General Transfer Format (GTF) between assemblies of the same, or closely-related species. Unlike current coordinate lift-over tools which require a pre-generated “chain” file as input, Liftoff is a standalone tool that takes two genome assemblies and a reference annotation as input and outputs an annotation of the target genome. Liftoff uses Minimap2 (Li, 2018) to align the gene sequences from a reference genome to the target genome. Rather than aligning whole genomes, aligning only the gene sequences allows genes to be lifted over even if there are many structural differences between the two genomes. For each gene, Liftoff finds the alignments of the exons that maximize sequence identity while preserving the transcript and gene structure. If two genes incorrectly map to overlapping loci, Liftoff determines which gene is most-likely mis-mapped, and attempts to re-map it. Liftoff can also find additional gene copies present in the target assembly that are not annotated in the reference.

Previously, we have used Liftoff to map genes to a new Ashkenazi human reference genome (Shumate, Zimin *et al.*, 2020) and to an updated assembly of the bread wheat genome, *Triticum aestivum* (Alonge, Shumate *et al.*, 2020). Here, in addition to describing the algorithm itself, we present two more examples demonstrating the accuracy and versatility of Liftoff. First, we map genes between two versions of the human reference genome. Next, to demonstrate a cross-species lift over, we map protein-coding genes from the human reference genome to a chimpanzee genome assembly.

## Implementation

Liftoff is implemented as a python command-line tool. The main goal of Liftoff is to align gene features from a reference genome to a target genome and use the alignment(s) to optimally convert the coordinates of each exon. An optimal mapping is one in which the sequence identity is maximized while maintaining the integrity of each exon, transcript, and gene. While our discussion of Liftoff here focuses on lifting over genes, transcripts, and exons, it will work for any feature, or group of hierarchical features present in a GFF or GTF file.

As input, Liftoff takes a reference genome sequence and a target genome sequence in FASTA format, and a reference genome annotation in GFF or GTF format. The reference annotation is processed with gffutils (https://github.com/daler/gffutils), which uses a sqlite3 database to track the hierarchical relationships within groups of features (e.g. gene, transcript, exon). Using pyfaidx (Shirley *et al.*, 2015), Liftoff extracts gene sequences from the reference genome, and then invokes Minimap2 to align the entire gene sequence including exons and introns to the target. The Minimap2 parameters are set to output up to 50 secondary alignments for each sequence in SAM format. By default, genes are aligned to the entire target genome, but for chromosome-scale assemblies, the user can enable an option to align genes chromosome by chromosome. Under that option, only those genes which fail to map to their expected chromosome are then aligned to the entire genome.

In many cases, a gene has a single complete alignment to the target genome, which makes finding the optimal mapping trivial. In other cases, differences between the two genomes cause the gene to align in many fragmented pieces, and the optimal mapping is some combination of alignments. To find this combination, Liftoff uses networkx (https://github.com/networkx/networkx) to build a directed acyclic graph representing the alignments as follows. Using Pysam (https://github.com/pysam-developers/pysam) to parse the Minimap2 alignments, each alignment is split at every insertion and deletion in order to form a group of gapless alignment blocks. Blocks not containing any part of an exon are discarded, and the remaining blocks are represented by nodes in the graph. Two nodes *u* and *v* are connected by an edge if the following conditions are true.

1. *u* and *v* are on the same chromosome or contig
2. *u* and *v* are on the same strand
3. *u* and *v* are in the correct 5’ to 3’ order
4. The distance from the start of *u* to the end of *v* in the target genome is no greater than 2 times that in the reference genome

Each node is assigned a weight equal to the number of mismatches within exons (mismatches in introns are not counted), and each edge is assigned a weight equal to the length of gaps within exons spanned by that edge. A source and sink are added to the graph representing the start and end of the gene respectively, and the shortest path from source to sink is found using Dijkstra's algorithm (Dijkstra and Others, 1959) where the weight function between two nodes *u* and *v* is

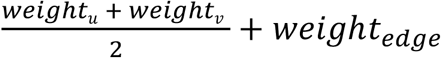

The shortest path represents the combination of aligned blocks that is concordant with the original structure of the gene and minimizes the number of mismatches and indels within exons. The alignments in this path define the final placement of the gene. Using the coordinates of the aligned blocks in the shortest path, the coordinates of each exon are converted to their respective coordinates in the target genome. A simple example of this process is shown in **Figure 1**, which illustrates lifting over a 5-exon transcript from the human reference genome (GRCh38) to a chimpanzee genome (PTRv2). This gene has a large intronic deletion in PTRv2 and does not have and end-to-end alignment, but it can still be successfully lifted over using our algorithm.

**Figure 1.**
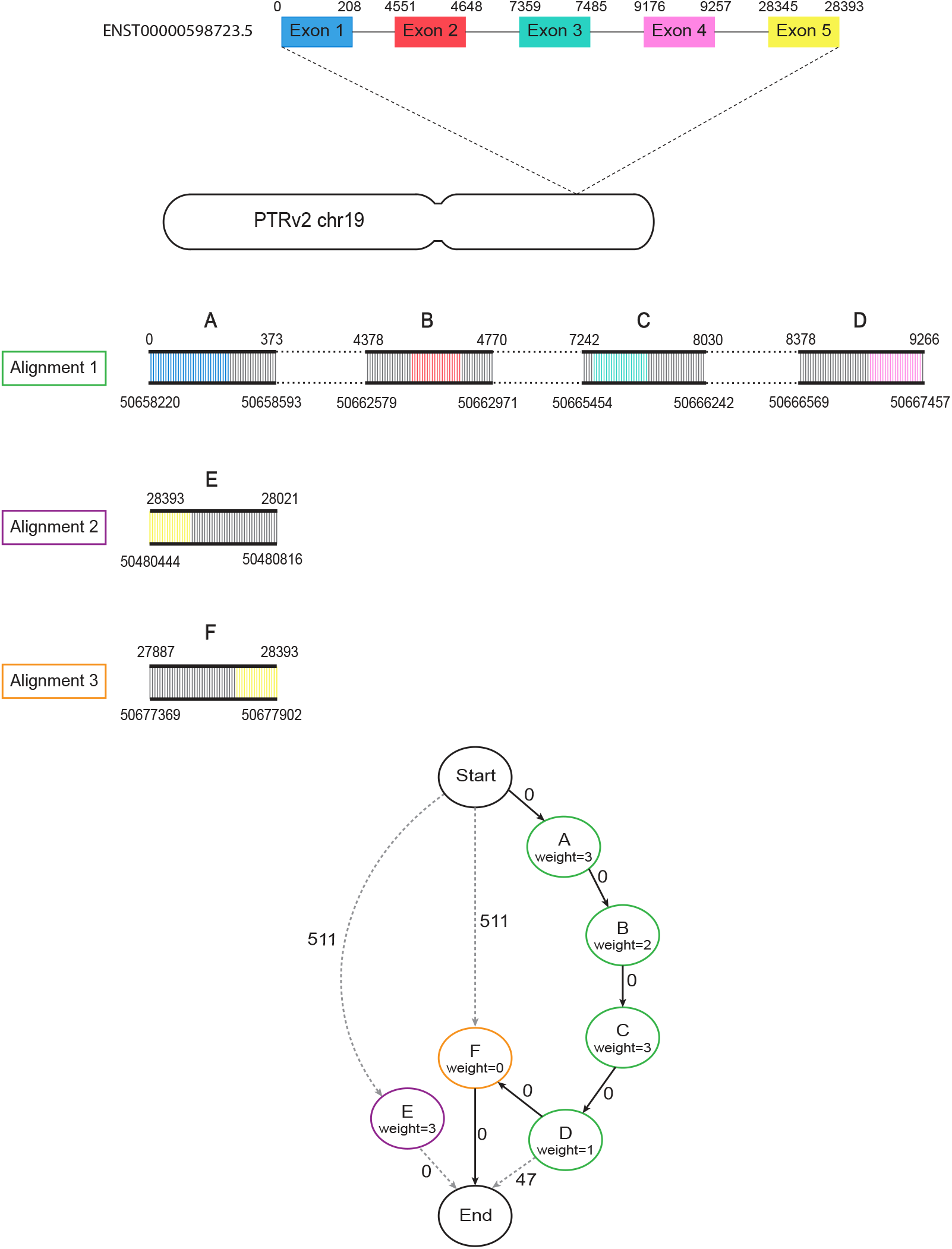
Example of the lift-over process. Diagram showing the steps taken by Liftoff when mapping human transcript ENST00000598723.5 to the chimpanzee (PTRv2) homolog on chromosome 19. Minimap2 produces 3 partial alignments of this gene to PTRv2. Alignment 1 (green) has 4 gapless blocks containing exons 1-4 which are represented by nodes A-D in the graph. The dashed lines in between blocks of the alignment represent gaps/introns. Alignments 2 (purple) and 3 (orange) each have 1 gapless block containing exon 5 represented by nodes E and F respectively. Node E is not on the same strand as alignments 1 and 2 and is therefore only connected to the start and end. The node weights correspond to the number of mismatches in exons and the edge weights are the number of unaligned exon bases between two nodes. The shortest path (A,B,C,D,F) is shown with bold arrows and contains complete alignments of all 5 exons with a total of 9 mismatches and 0 gaps.

One of the main challenges with gene annotation lift over is correctly mapping homologous genes from multi-gene families. Two different genes may optimally map to the same locus if they are identical or nearly identical. To handle this situation, after Liftoff maps all genes to their best matches, it checks for pairs of genes on the reference genome that have incorrectly mapped to overlapping (or identical) locations on the target genome, and it then attempts to find another valid mapping for one of the genes. Liftoff first tries to remap the gene with the lower sequence identity. If the genes mapped with the same sequence identity, Liftoff considers the neighboring genes and tries to remap the gene that appears out of order according to the reference annotation. When remapping the gene, Liftoff rebuilds the graph of aligned blocks excluding any blocks that overlap the homologous gene. The shortest path through this new graph represents the best mapping for this gene that does not overlap its homolog. If another valid mapping does not exist, the gene with lower identity is considered unmapped. This process is repeated until there are no genes mapped to overlapping loci. Liftoff then outputs a GFF file with the coordinates on the target genome of all of the features from the original annotation, and a text file with the IDs of any genes that could not be lifted over.

Note that differences in the genome sequences themselves may result in Liftoff mapping a gene to a paralogous location. For example, consider a gene family with 5 members on the reference genome but only 4 members on the target. The fifth gene might simply be unmapped, but if the target has a paralogous copy elsewhere, *and* if that copy is not matched by a homolog on the reference, then Liftoff will map the fifth gene to the paralogous location.

## Annotating Extra Gene Copies

Another feature unique to Liftoff is the option to find additional copies of genes in the target assembly not annotated in the reference. With this option enabled, Liftoff maps the complete reference annotation first, and then repeats the lift-over process for all genes. An extra gene copy is annotated if another mapping is found that does not overlap any previously-annotated genes, and that meets the user-defined minimum sequence identity threshold. The lift-over procedure is repeated until all valid mappings have been found.

We recently used Liftoff with this feature enabled to annotate our improved assembly of the bread wheat genome, which contains 15.07 gigabases of anchored sequence compared to 13.84 in a previous reference genome (Alonge, Shumate *et al.*, 2020). In addition to successfully mapping 100,839 of the 105,200 reference genes to this large and complex genome, we found 5,799 additional gene copies using a strict sequence identity threshold of 100%.

## Results

Here we demonstrate Liftoff’s ability to lift an annotation to an updated reference genome by lifting genes from the two most recent versions of the human reference genome, GRCh37 and GRCh38. We also demonstrate Liftoff’s ability to lift genes between genomes of closely-related species by lifting genes from GRCh38 to the chimpanzee genome Clint_PTRv2. To assess the accuracy of Liftoff in each example, we evaluate both the sequence identity and order of mapped genes.

### GRCh37 to GRCh38

We attempted to map all protein-coding genes and lncRNAs on primary chromosomes in the GENCODE v19 annotation (Harrow *et al.*, 2012) from GRCh37 to GRCh38. Out of 27,459 genes, we successfully mapped 27,424 (99.87%). We consider a gene to be successfully mapped if at least 50% of the reference gene maps to the target assembly. An overwhelming majority of the gene sequences in GRCh38 were nearly identical to the sequences in GRCh37, with an average sequence identity in exons of 99.97% (**Figure 2**).

**Figure 2.**
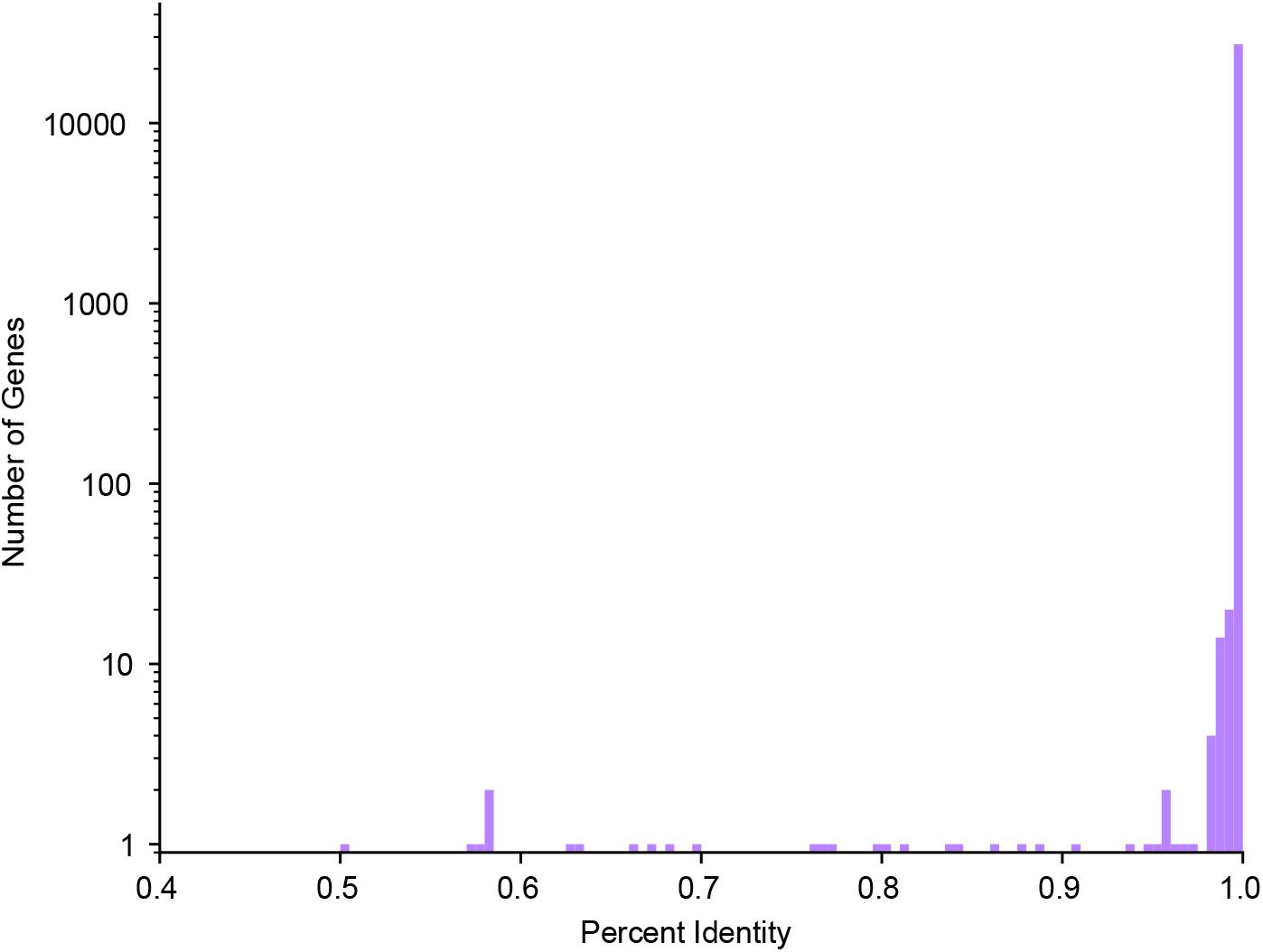
Distribution of GRCh37 and GRCh38 sequence identity. Histogram showing the distribution of exon sequence identity of protein-coding and lncRNA genes in GRCh37 and GRCh38. Log scale used to make the counts of just 1 or 2 genes visible; all bins below 97% identity contain at most 2 genes.

To visualize the co-linearity of the gene order between the two assemblies, we plotted each gene as a single point on a 2D plot where the X coordinate is the ordinal position of the gene in GRCh37 and the Y coordinate is the ordinal position in GRCh38 (**Figure 3**).

**Figure 3.**
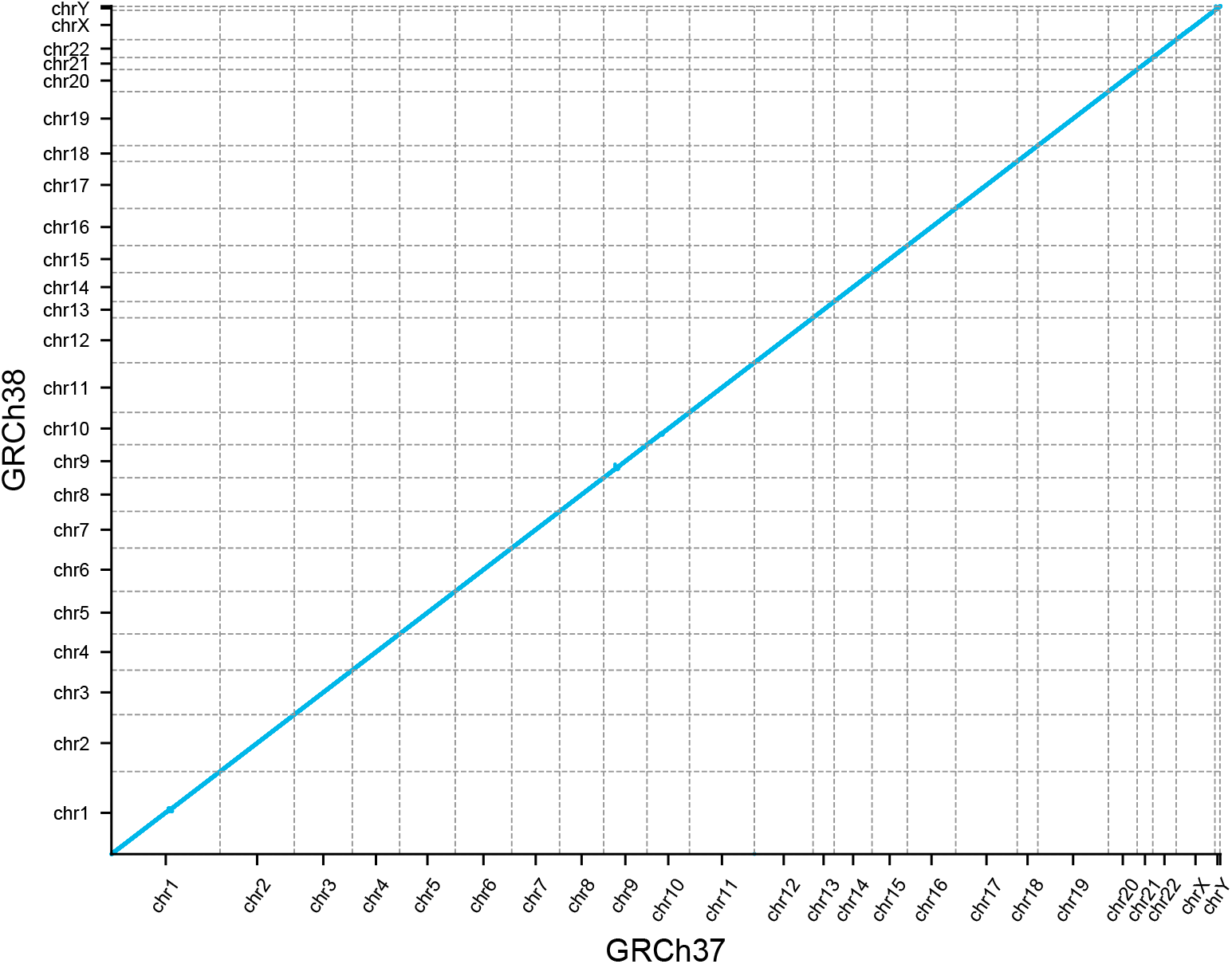
GRCh37 and GRCh38 gene order. Dot plot showing the ordinal position of each gene in GRCh37 on the x-axis and the ordinal position in GRCh38 on the y-axis.

The gene order appears perfectly co-linear; however, there are some exceptions not visible at the scale of the whole genome. To calculate the number of genes out of order in GRCh38 with respect to GRCh37, we first sorted the X,Y points by X, and then found the length of the longest increasing subsequence in Y. The longest increasing subsequence in Y represents the genes in GRCh38 that are in order with respect to GRCh37, and those points not belonging to this subsequence are genes which are out of order. With this process we found 305 genes (1.1%) occurring in a different relative position in GRCh38 with respect to GRCh37.

### GRCh38 to PTRv2

We attempted to map all protein-coding genes on chromosomes 1-22 and chromosome X in the GENCODE v33 annotation (Frankish *et al.*, 2019) from GRCh38 to an assembly of the chimpanzee (*Pan troglodytes*), PTRv2. Out of 19,878 genes, we were able to map 19,555 (98.38%). The average sequence identity in exons of successfully mapped genes was 98.70% (**Figure 4**).

**Figure 4.**
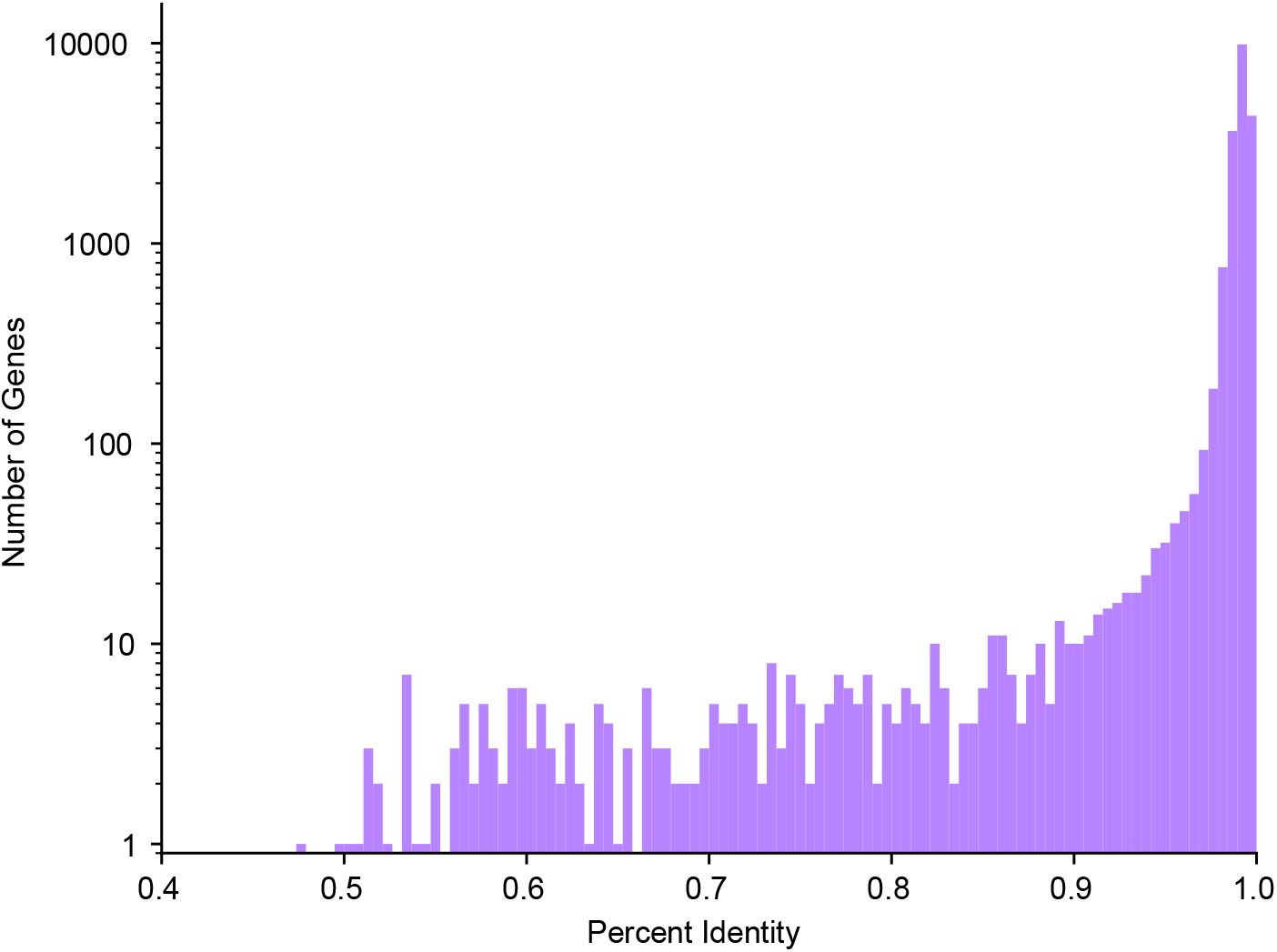
Distribution of GRCh38 and PTRv2 sequence identity. Histogram showing the distribution of exon sequence identity of protein-coding genes in GRCh38 and PTRv2. Note that the y-axis is shown on a log scale, as in Figure 2.

As was done with the GRCh37 to GRCh38 lift-over, we compared the gene order in GRCh38 to that in PTRv2 and found 2,172 genes in PTRv2 to be in a different relative position. Some of these ordinal differences are visible at the whole-genome scale (**Figure 5**) including 4 large regions on the chimpanzee homologues of chromosomes 4, 5, 12, and 17 where the gene order is inverted due to large-scale chromosomal inversions.

**Figure 5.**
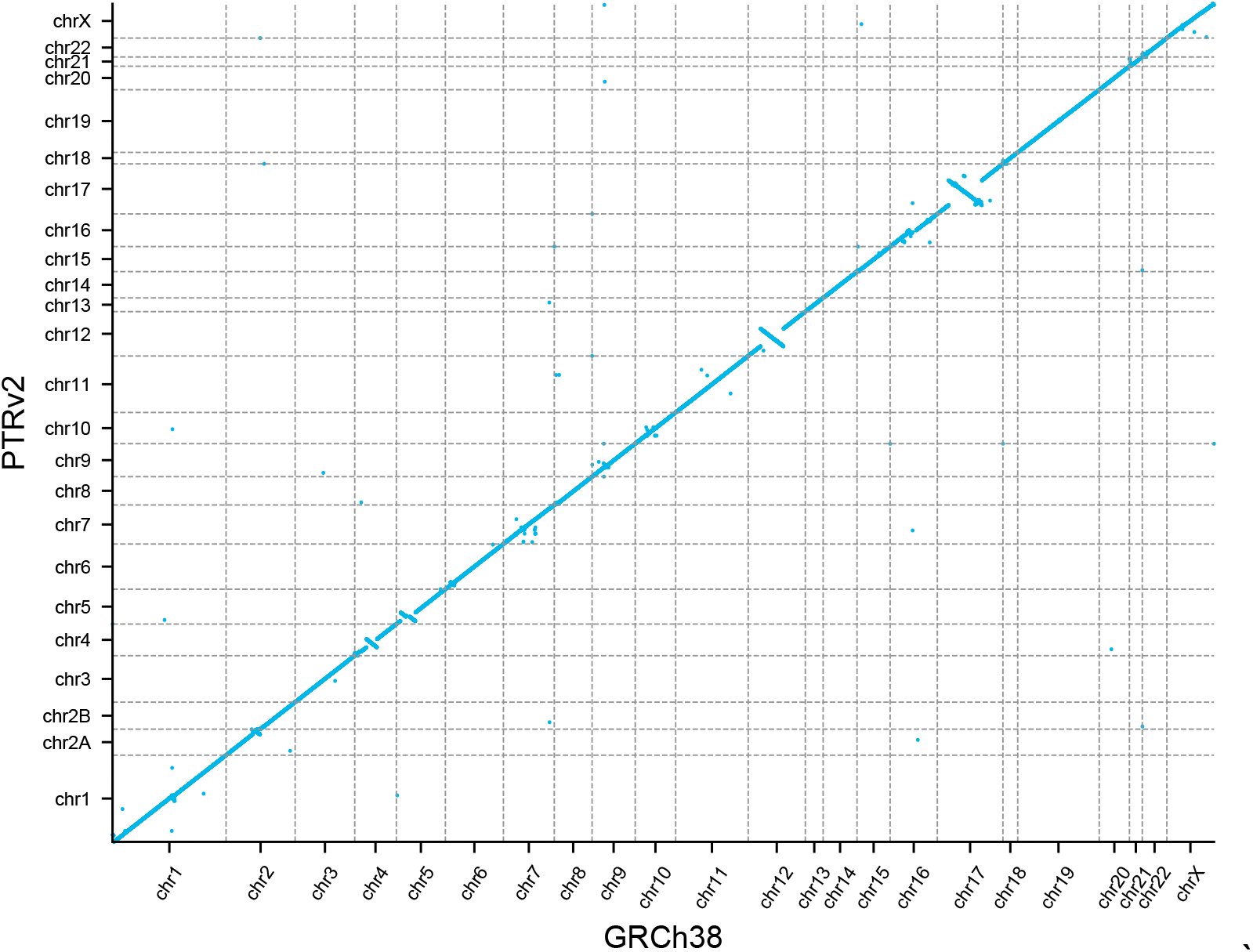
GRCh38 and PTRv2 gene order. Dot plot showing the ordinal position of each gene in GRCh38 on the x-axis and the ordinal position in PTRv2 on the y-axis.

## Discussion

The rapidly growing number of high-quality genome assemblies has greatly increased our potential to understand sequence diversity, but accurate genome annotation is needed to understand the biological impact of this diversity. Rather than annotating genomes *de novo*, we can take advantage of the extensive work that has gone into creating reference annotations for many well-studied species. We developed Liftoff as an accurate tool for transferring gene annotations between genomes of the same or closely-related species. Unlike current coordinate lift-over strategies which only consider sequence homology, Liftoff considers the constraints between exons of the same gene and constraint that distinct genes need to map to distinct locations.

We showed that we were able to lift over nearly all genes from GRCh37 to GRCh38. The gene sequences and order are very similar between the two assemblies, with an average sequence identity of >99.9% and only 305 genes appearing in a different order. GRCh38 fixed a number of mis-assemblies and single base errors present in GRCh37 (Guo *et al.*, 2017), so it is not unexpected that the gene sequence and order are not entirely identical. This demonstrates Liftoff’s ability to accurately annotate an updated reference assembly, making it a useful tool as reference assemblies are continuously updated.

We also showed that we could lift-over nearly all protein-coding genes from GRCh38 to the chimpanzee genome, PTRv2, with an average sequence identity of 98.7%. This is consistent with previous work showing the human genome and chimpanzee genome are approximately 98% identical (Chimpanzee Sequencing and Analysis Consortium, 2005). Comparing the gene order revealed 4 large regions on the homologs of chromosomes 4, 5, 12, and 17 where the gene order is inverted. These regions are consistent with previous reports: the chimpanzee genome has 9 well-characterized pericentric inversions on chromosome homologs 1, 4, 5, 9, 12, 15, 16, 17 (Yunis and Prakash, 1982). The 4 largest of these inversions are on 4, 5, 12, and 17 (Soto *et al.*, 2020) hence their visibility at this scale. Additionally, the co-linear mapping of genes from human chromosome 2 to chimpanzee chromosomes 2A and 2B is consistent with the known telomeric fusion of these chromosomes (Yunis and Prakash, 1982). The consistency of the gene sequence identity with the known genome sequence identity between chimpanzee and human, and the consistency of the gene order with the known structural differences between the two genomes demonstrate the accuracy of Liftoff’s gene placements in a cross-species lift-over.

Annotating new assemblies with a lift-over strategy rather than *de novo* is limited in that the annotation of the new assembly will only be as complete as the reference. However, as reference annotations continue to improve through manual curation, experimental validation or improved computational methods, Liftoff will enable easy integration of these improvements across many genomes. We anticipate that Liftoff will be a valuable tool in improving our understanding of the biological function of the large and rapidly growing number of sequenced genomes.

## Acknowledgements

This work was supported in part by NIH under grants R01-HG006677 and R35-GM130151.

